# Environmental DNA persistence and fish detection in captive sponges

**DOI:** 10.1101/2022.04.20.488927

**Authors:** Wang Cai, Lynsey R. Harper, Erika F. Neave, Peter Shum, Jamie Craggs, María Belén Arias, Ana Riesgo, Stefano Mariani

## Abstract

Large and hyper-diverse marine ecosystems pose significant challenges to biodiversity monitoring. While environmental DNA (eDNA) promises to meet many of these challenges, recent studies suggested that sponges, as ‘natural samplers’ of eDNA, could further streamline the workflow for detecting marine vertebrates. However, beyond pilot studies demonstrating the ability of sponges to capture eDNA, little is known about the journey of eDNA particles in the sponge tissues, and the effectiveness of the latter compared to water samples. Here, we present the results of a controlled aquarium experiment to examine the persistence and detectability of eDNA from three encrusting sponge species and how these compare with established water filtration techniques. Our results indicate that sponges and water samples have highly similar detectability for fish of different sizes and abundances, but different sponge species exhibit considerable variance in performance. Interestingly, one sponge appeared to mirror the eDNA degradation profile of water samples, while another sponge retained eDNA throughout the experiment. A third sponge yielded virtually no DNA sequences at all. Overall, our study suggests that some sponges will be suitable as natural samplers, while others will introduce significant problems for laboratory processing. We suggest that an initial optimization phase will be required in any future studies aiming to employ sponges for biodiversity assessment. With time, factoring in technical and natural accessibility, it is expected that specific sponge taxa may become the ‘chosen’ natural samplers in certain habitats and regions.

## Introduction

The advancing method of environmental DNA (eDNA) analysis is increasingly used to profile biodiversity in ecological research (Bohmann et al., 2014). This method can target multiple taxa in parallel by capturing, extracting and sequencing DNA from exfoliated cells and extracellular DNA from different environmental samples, such as water, soil, and air (Andersen et al., 2012; Eble et al., 2020; Lynggaard et al., 2022), followed by taxonomic assignment using bioinformatic tools (Cristescu et al., 2014). High-throughput capability and low requirements for on-site taxonomists mean that eDNA analysis offers considerable improvements over certain traditional survey methods (Lebuhn et al., 2013; Goldberg et al., 2016), thus research on and applications of eDNA have proliferated in the past decade (Pawlowski et al., 2020). To date, eDNA has been most extensively used to monitor aquatic biodiversity and address important ecological questions in aquatic systems (Ruppert et al., 2019).

Environmental DNA is especially beneficial to marine research, where the deployment of large-scale and multi-taxa biodiversity surveys is challenging and costly. In the marine environment, studies have typically compared eDNA performance with well-established catch-based and video-based methods (Russo et al., 2021; Valdivia-Carrillo et al., 2021). These studies generally show that eDNA is an effective and sensitive method for marine biodiversity assessment, often outperforming the traditional approaches (Boussarie et al., 2018; Aglieri et al., 2021). Nevertheless, existing marine eDNA protocols are not without challenges, given the sheer size and considerable physical and ecological complexity of marine environments (Hansen et al., 2018). One limitation of marine eDNA is the sampling capability (Goldberg et al., 2016).

Aquatic eDNA is primarily collected from the sea using water filtration via an artificial membrane (McQuillan et al., 2017). Unsurprisingly, researchers advised that large volume filtration and increased sampling replication should be considered in eDNA studies to avoid false negatives (Bessey et al., 2020; Stauffer et al., 2021). However, these optimizations require a substantial budget for study design (Ficetola et al., 2015) or lead to significant investment in high-tech solutions such as integrated eDNA sampling systems (Thomas et al., 2018) and deep-sea robotic samplers (McQuillan et al., 2017). These high-tech solutions can become limiting for small research groups in many parts of the world and studies in remote and poorly accessible environments. As an alternative to mechanical filtration systems, passive eDNA sampling has been proposed to lower technological investment via simple capture media. For example, Kirtane et al. (2020) and Bessey et al. (2021a) conducted and tested adsorbent-filled sachets to collect and preserve eDNA, while Bessey et al. (2021b) examined the efficiency of submerging filter membranes directly in the water column, thereby eliminating labour intensive water filtration. Another line of research, which further reduces deployment times and the use of gear and plastics, is the harnessing of natural eDNA samplers, i.e., live filter-feeding organisms in aquatic ecosystems. Siegenthaler et al. (2019) used gut contents from shrimps to assess fish diversity, and Wells et al. (2021) used anemones’ diet to assess plankton communities. Mariani et al. (2019) found that eDNA extracted from sponges can detect the presence of a variety of marine fish and mammals in Mediterranean and Antarctic waters. In addition, Turon et al. (2020) further highlighted the usefulness of sponges to describe tropical fish communities from coral reefs in South-eastern Asia.

This new perspective opens up possibilities for relatively low-cost and low-tech biodiversity monitoring. Sponges are the most efficient natural water filters on the planet (Kahn et al., 2015), and their pumping rates can vary from 0.3 to 35 ml/min /cm^3^ (Gerrodette & Flechsing, 1979; Hoffmann et al., 2008; Weisz et al., 2008). Thanks to their structure, sponges can trap objects ranging in size from microscopic particles to relatively large diatoms (Ribes et al., 1999; Riesgo et al., 2021), and have been used to uncover sponge-associated Arthropoda and Annelida communities (Kandler et al., 2019). Given their ubiquitous distribution (Van Soest et al., 2012), regeneration ability (Ereskovsky et al., 2021), and ease of sampling, sponges have the potential to become cost-effective natural samplers in marine ecosystems for eDNA surveys. However, sponge eDNA pumping/trapping ability is likely to be affected by many factors, such as size (Morganti et al., 2019) and symbiont content (Weisz et al., 2008), which may contribute to whether certain sponge species can serve as natural eDNA samplers.

To our knowledge, no direct comparisons of eDNA capture from sponge and water samples have been carried out yet in controlled or natural settings. Here, we designed a tank experiment to compare the persistence and detectability of eDNA between three sponge species and standard water filtering protocols. We introduced fish species in replicated tanks with sponges for 40 hours to allow sponges to accumulate eDNA, then removed the fish. Repeat sampling over a period of four days offered insights into fish eDNA degradation and detectability variance between sponges and water samples, with important implications for the future use of sponges as natural eDNA samplers.

## Materials and Methods

### Experimental design

#### Experimental facilities and materials

Aquaria experiments were completed at the Horniman Museum & Gardens (HMG), London. We set up three independent aquarium systems, each comprising a 165L experimental tank and an 80L filtration sump located underneath. This sump contained mechanical filtration (ClariSea SK5000 Generation 2), 3.5kg of biological filtration (Maxspect Nano Tech Bio Spheres), and a protein skimmer (Bubble Magus Curve 9). A return pump located in the sump continuously supplied high-quality, temperature-controlled (T_mean_ = 26.8°C) seawater to the experimental tank. Water then returned directly into the mechanical filtration of the sump via a 40mm standpipe. Our experiments employed three sponge species: *Chondrilla* sp. (order Chondrosiida), *Axinyssa* sp. (order Suberitida), and *Darwinella* sp. (order Dendroceratida). These sponges were chosen as they had naturally colonised the coral colonies at the HMG aquarium facilities, and because of their different filtering characteristics. *Chondrilla* sp. has a slower pumping rate than *Axinyssa* sp. and *Darwinella* sp., and is known as a high microbial abundance (HMA) sponge type (Moitinho-Silva et al. 2017; Batista et al. 2018; Díez-Vives et al. 2020). These colonising sponges at the aquarium facilities offered an ideal head start for our comparative experiment, as it is typically difficult to *de-novo* rear sponges with different filtration characteristics simultaneously in artificial environments.

Before being placed into the experimental tanks, the sponges were quarantined in a separate empty tank (with no fish) for seven days to ensure that fish eDNA would be removed from their tissue prior to the subsequent experiments.

Five fish species were employed in replicated tanks. Three individuals of clown anemonefish (*Amphiprion ocellaris*) and three individuals of blue-green damselfish (*Chromis viridis*) were placed into each tank. We placed one individual of sailfin tang (*Zebrasoma veliferum*) into tank A, one individual of royal gramma (*Gramma loreto*) into tank B, and one individual of half-spined seahorse (*Hippocampus semispinosus*) into tank C. Each tank contained the same three species of sponges, two fish species common across the three tanks, and one fish species unique to each tank (Fig. 1).

**Figure 1.**
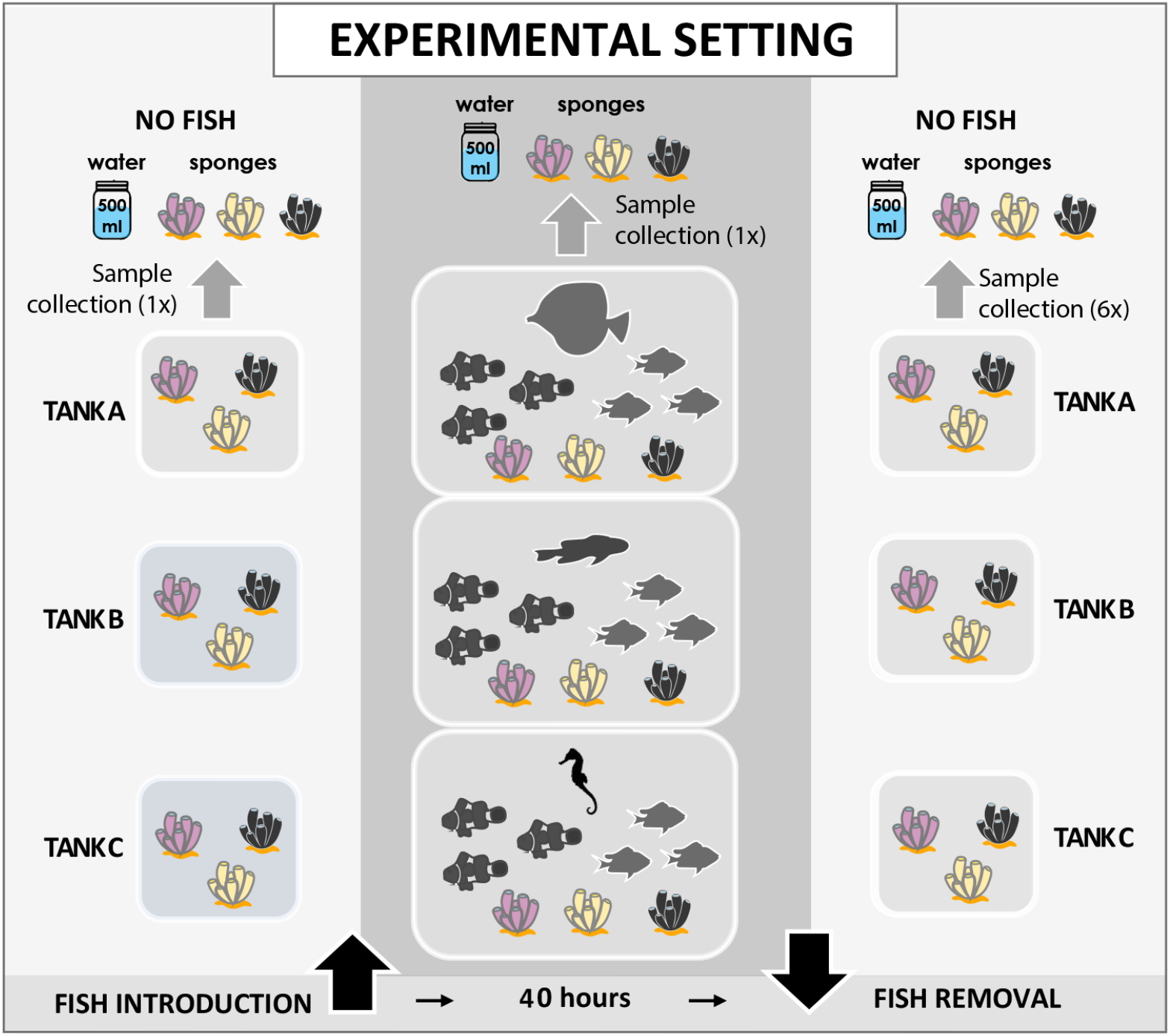
Experimental setting. There are three main phases of the experiment. The first phase took place prior to introducing fish into tanks, when there were only sponges. The second phase involved placing fish into the experimental tanks for 40 hours. In the last phase, fish were removed and sponges remained in tanks for 72 hours. Samples were collected at six time points: 0, 4, 8, 24, 48, and 72 hours after removing fish in the last phase. A 500 mL water sample and one biopsy of each sponge species were collected at each time point for each tank. The water samples were filtered using a syringe with a sterivex filter, while sponge biopsies were preserved in 100% ethanol.

#### DNA degradation experiment

Sponges were allowed to acclimate to experimental tanks for three days. Fish were introduced and remained in the tanks unfed for 40 hours, then all fish were removed while sponges remained in the tanks for another 72 hours. Water and sponge samples were collected at eight time points (Fig. 1): one sample was taken before fish introduction (considering the possibility that sponges still contained or introduced eDNA from non-experimental fish), one sample was taken when fish had been in the tanks for 20 hours (considering the possibilities that eDNA may already be present in the water column and sponge tissue or that eDNA may not yet be detectable in water or sponges), then samples were taken at 0, 4, 8, 24, 48, and 72 hours after fish removal.

For each time point in each tank, we collected one water sample and three sponge samples (one biopsy for each species), totalling 96 samples (4 samples × 8 time points × 3 tanks). A 500 mL water sample was collected and filtered onto a 0.45 μm Sterivex filter (PES membrane, Merck Millipore) using a 60 mL syringe (Fisher Scientific). Each filter was placed into a single bag and then immediately stored at −20°C until DNA extraction. Using arm-length gloves with wrist gloves on top, sponge biopsies were taken using disposable scalpels (Swann-Morton No 21) and stored in 2 mL tubes with 100% ethanol at −20°C. At the end of the experiment, all sponge biopsies were transferred into fresh 2 mL tubes with 100% ethanol and stored at −20°C until DNA extraction. The sponge biopsies sampled were much smaller than the whole individual (around 1 cm^3^, ≤25 mg) to minimise stress on the organism.

In addition, we included one seawater blank from the filtration system (500 mL), one filtration blank (500 mL of MilliQ water), and one sponge blank (artificial kitchen sponge submerged in a sterile 10 L plastic box containing seawater from the filtration system) on each experimental day to assess for potential contamination. All sampling implements (water bottle, tubes, scalpel, long glove, and laboratory materials) were sterilized prior to and disposed of after sample collection. Between sampling events, workspaces were decontaminated using 10% v/v bleach solution followed by 70% v/v ethanol solution.

### Laboratory procedures

Each filter capsule was opened onto a petri dish using carpenter pliers. The filters were removed from the inner tube and torn into small pieces using metal forceps. All pieces were placed inside a 1.5 mL microtube for DNA extraction using the Mu-DNA water protocol (Sellers et al. 2018). For sponge biopsies, approximately 25 mg of dry sponge per sample was used for DNA extraction. Each sponge was removed from the storage ethanol and blotted dry against filter paper (42.5 mm, Fisher Scientific) inside a petri dish, then processed using the Mu-DNA tissue protocol with an inhibitor removal step the Mu-DNA water protocol (Sellers et al. 2018). The extracts were then quantified for DNA concentration using a Qubit™ 4 fluorometer with a Qubit™ dsDNA HS Assay kit (Thermo Fisher Scientific). Sponge DNA extracts were diluted 1:10 if initial PCRs failed to enable PCR amplification for library preparation.

All reusable equipment for DNA extraction was first sterilised in 10% v/v bleach solution, followed by a rinse in 5% v/v Lipsol detergent and deionised water. Equipment and consumables were then exposed to 30 minutes of ultraviolet (UV) light.

PCR amplifications were carried out using the Tele02 primers (Taberlet et al. 2018), which amplify a ∼169 bp fragment of the mitochondrial 12S rRNA gene. Primers contained unique 8-bp dual barcodes for sample identification and to reduce tag jumping (Schnell et al., 2015), with 2-4 leading ‘N’ bases to increase sequence diversity. Samples were amplified in triplicate under the following conditions: initial denaturation at 95°C for 10 min, followed by 40 cycles of 95°C for 30 s, 60°C for 45 s, 72°C for 30 s, and finishing at 72°C for 5 mins. All PCRs were performed in 20 μl reactions containing 10 μl 2X MyFi Mix (Meridian Bioscience), 0.5 μM of each primer, 0.04 mg BSA (Bovine Serum Albumin Solution, Thermo Fisher Scientific), 5.84 μl of molecular grade water (Invitrogen), and 2 μl DNA template. Two or three PCR positive/negative controls were included on each PCR run for eDNA samples and nsDNA samples (higher number of nsDNA samples than eDNA samples), respectively. PCR positive controls (0.05 ng/μl) contained one fish species which was not present in the HMG (the iridescent catfish *Pangasionodon hypopthalmus*). PCR triplicates were pooled together and visualized on 2% agarose gels. Each sample was purified using Mag-Bind Total Pure NGS (Omega Bio-Tek) magnetic beads. Purified PCR products were quantified as above and pooled in equimolar amounts to create two libraries, one for eDNA samples (24 samples and 18 controls), and one for nsDNA samples (72 samples and 16 controls). Pooled PCR products were purified with magnetic beads, and libraries were prepared using the NEXTFLEX Rapid DNA-Seq Kit for Illumina (PerkinElmer) following the manufacturer’s instructions. The libraries were quantified by qPCR using the NEBNext Library Quant Kit for Illumina (New England Biolabs) and Tape Station 4200 (Agilent). Then they were pooled in equimolar concentrations with a final molarity of 60 pM with 10% PhiX control. The libraries were sequenced on an Illumina iSeq100 using iSeq i1 Reagent v2 (300 cycles) at Liverpool John Moores University. This sequencing run included six samples from other projects but were removed prior to bioinformatic processing.

### Bioinformatic and Statistical analysis

The bioinformatic processing was based on the OBITOOLS software 1.2.11 (Boyer et al. 2016). The raw sequencing data were first trimmed to remove low-quality ends using ‘obicut’. After trimming, paired-end reads were merged by ‘illuminapairedend’, and alignments with low (<40) quality scores were removed. The alignments were then demultiplexed using ‘ngsfilter’ with default parameters. Subsequently, quality filters were performed by ‘obigrep’ to retain sequences between 130 bp to 190 bp without ambiguity to filter out erroneous sequences, followed by dereplication using ‘obiuniq’, and chimera removal using the *de novo* chimera search function in vsearch 2.4.3 (Rognes et al. 2016). The remaining sequences were clustered by SWARM v2 (Mahé et al. 2015) with ‘-d 3’. Taxonomic assignment was performed via ‘ecotag’. The reference database used in ‘ecotag’ was constructed by *in silico* PCR for Tele02 primers against the EMBL database (Release version r143) using ‘ecoPCR’. An additional taxonomic assignment was carried out using BLAST against the NCBI reference database to check the assignment of sequences, and we manually corrected one species (*Gramma loreto*) that ‘ecotag’ only can assign to the metazoan level. Finally, a sample/OTU table with taxon information was formatted using R scripts listed at https://github.com/metabarpark/R_scripts_metabarpark. We then used the R package “lulu” 0.1.0 (Frøslev et al., 2017) with default parameters to filter erroneous OTUs based on the calculation of OTUs’ pairwise similarities and co-occurrence patterns. The remaining OTUs were further collapsed using the metabarpark ‘owi_collapse’ R script.

All downstream statistical analyses were carried out in R v3.6.3. The raw read counts were first normalized using a log_10_ transformation. We visualized community composition by the ‘pheatmap’ function (v1.0.12; Kolde, 2019). To examine the relationship between time and the total log_10_-reads of each sample, we performed linear models (LMs) using the ‘lm’ function. We then converted the log_10_ read counts to presence/absence (1/0) for subsequent analyses. To compare pairwise differences in species richness across samples, we performed a multiple pairwise test by the ‘TukeyHSD’ function. ‘TukeyHSD’ is a post-hoc test, which allows us to comparing the differences between two sample types based on analysis of variance. Subsequently, we used ‘mvabund’ v3.12.3 (Wang et al. 2012) to analyse the effects of covariates (sample type, time, and tank) on community composition. ‘mvabund’ is a model-based method (generalised linear model framework) which allows us to select an appropriate error distribution for the corresponding data type. Here we select binomial error distribution for the presence/absence data. All results were visualized using the ‘ggplot2’ R package v 3.3.5.

## Results

### Bioinformatic processing and taxonomic composition

Sequencing yielded nearly 1 million raw sequences, of which 703,816 for the sponge nsDNA library and 295,854 for the water eDNA library. After quality filtering, we retained 652,148 paired-end reads, clustered into 175 OTUs. OTUs were assigned to experimental fish species, PCR positive control species, human, non-experimental fish species present in the HMG facilities and in feeds. 119 OTUs (8.6% read counts) could not be identified below the Class level (Table S1). Three experimental fish species were assigned to species-level, and two were assigned to genus-level with high confidence (100%). Five OTUs were assigned to non-experimental fish species (genus-level, >97% identity). These non-experimental species were probably carried over to the experimental setup by the sponges, as most were detected at the beginning of the experiment (Table S1). We then chose a stringent level to minimize false positives and contamination by removing low-abundance read counts of OTUs (10 ≤ reads), according to highest read counts of non-positive control species and lowest read counts of true positives species in PCR positive control. After filtering, all blanks were clean, indicating that sampling and decontamination procedures were appropriate and false positives/contamination would not affect downstream analysis. In addition, the non-experimental fish OTUs detected in the corresponding tanks were removed based on the samples collected before fish introduction. The details of species composition are shown in Figures S2 and S3. For downstream analysis, we only considered the five experimental fish species.

The total read counts for the five experimental species decreased across samples over time (Fig. 2A) but varied among samples. There was a significant negative relationship between the total reads and time (p < 0.001, R^2^ = 0.443) for the aquatic eDNA samples and the sponge *Axinyssa* sp. (p = 0.009, R^2^ = 0.276), while the total reads of *Darwinella* sp. showed no correlation with time (p = 0.233, R^2^ = 0.025). *Chondrilla* sp. did not show any positive detection over time.

**Figure 2.**
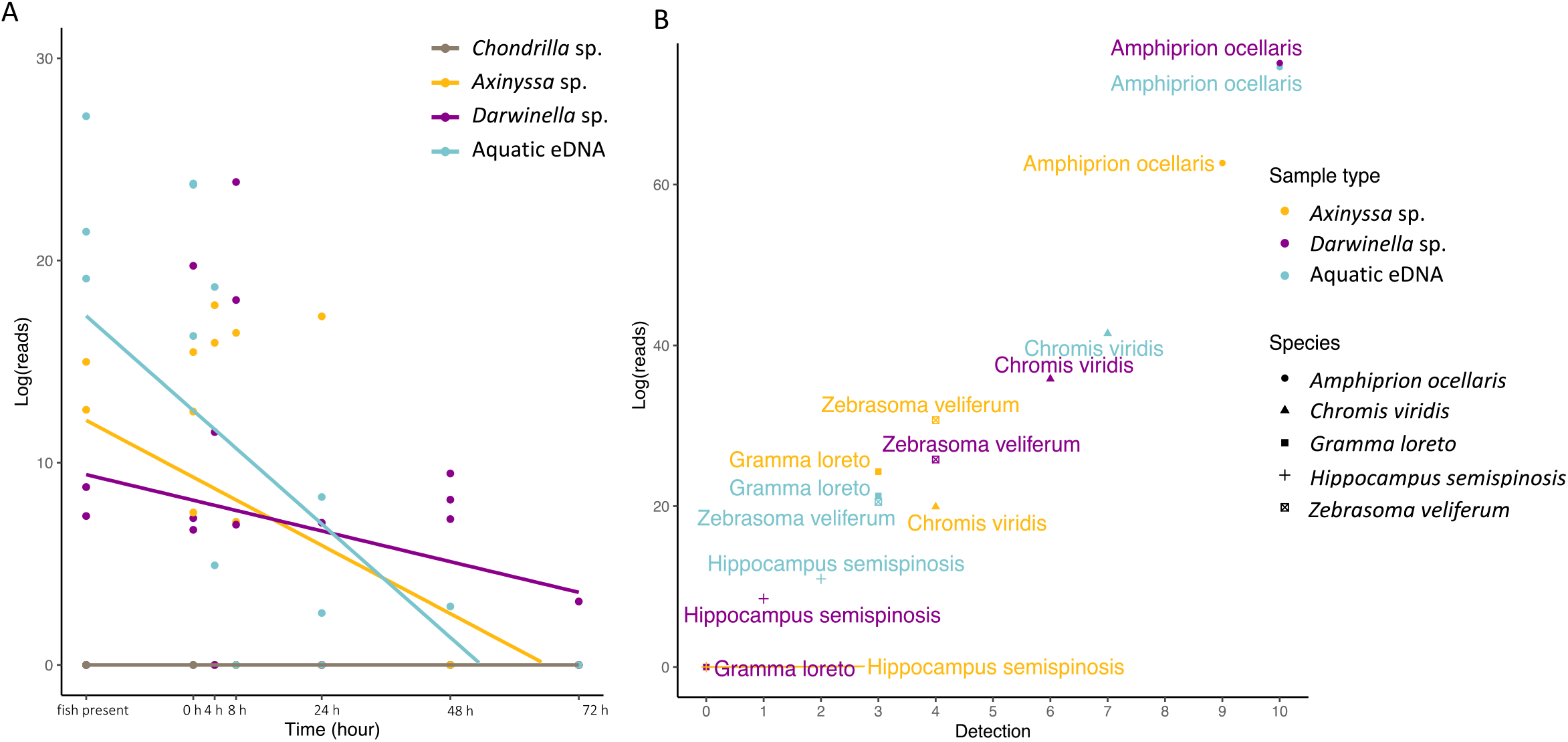
Comparing the total read counts of each sample and the detectability of each species. A. The linear relationship between the total log_10_ read counts and the time point of each sample type. The points represent sample replicates per tank. The x-axis is the timeline of the sampling events. The “fish present” represents fish were in the tanks, the time represents the time point after fish removal. **B**. The detectability of each fish species per sample type. Shapes represent species. The y-axis is the total log_10_ read counts of each species per sample type, and the x-axis is the total number of positive detections of each fish per sample type. The colours represent the sample type for both A and B.

### The detectability of each species across nsDNA and eDNA

To compare the detection efficiency of different fish between sponge nsDNA and aquatic eDNA, we examined the detection rate and log reads per species for each sample (Fig. 2B). Overall, detection frequency was correlated with read abundance, but varied considerably among the experimental species. The abundant species *Amphiprion ocellaris* and *Chromis viridis* (three individuals in each tank) had higher detection rates than *Z. veliferum, G. loreto* and *H. semispinosus* (one individual in each tank), however, *A. ocellaris* and *C. viridis* also differed from each other. Furthermore, the relatively larger individual (*Z. veliferum*) had a higher detection rate than the other two smaller, unique fishes. This pattern was consistent across aquatic eDNA and sponge nsDNA samples (Fig. 2B).

### Degradation of nsDNA and eDNA

The linear model analysis found significantly different decay patterns between sponge nsDNA and aquatic eDNA (Fig. 3). We used presence/absence data, focusing on species richness rather than read abundance. Species richness declined steeply over time in water samples (p < 0.001, R^2^ = 0.437), and also significantly, albeit less steeply, in the sponge *Axinyssa* sp. (p = 0.008, R^2^ = 0.278), while *Darwinella* sp. showed no significant decrease over the duration of the experiment (p = 0.424, R^2^ = −0.017, Fig. 3B). Time effect aside, there was no significant difference in species richness between aquatic eDNA and sponge nsDNA during the observation period (Tukey’s comparisons, p > 0.5, omitting *Chondrilla* sp., which failed to amplify).

**Figure 3.**
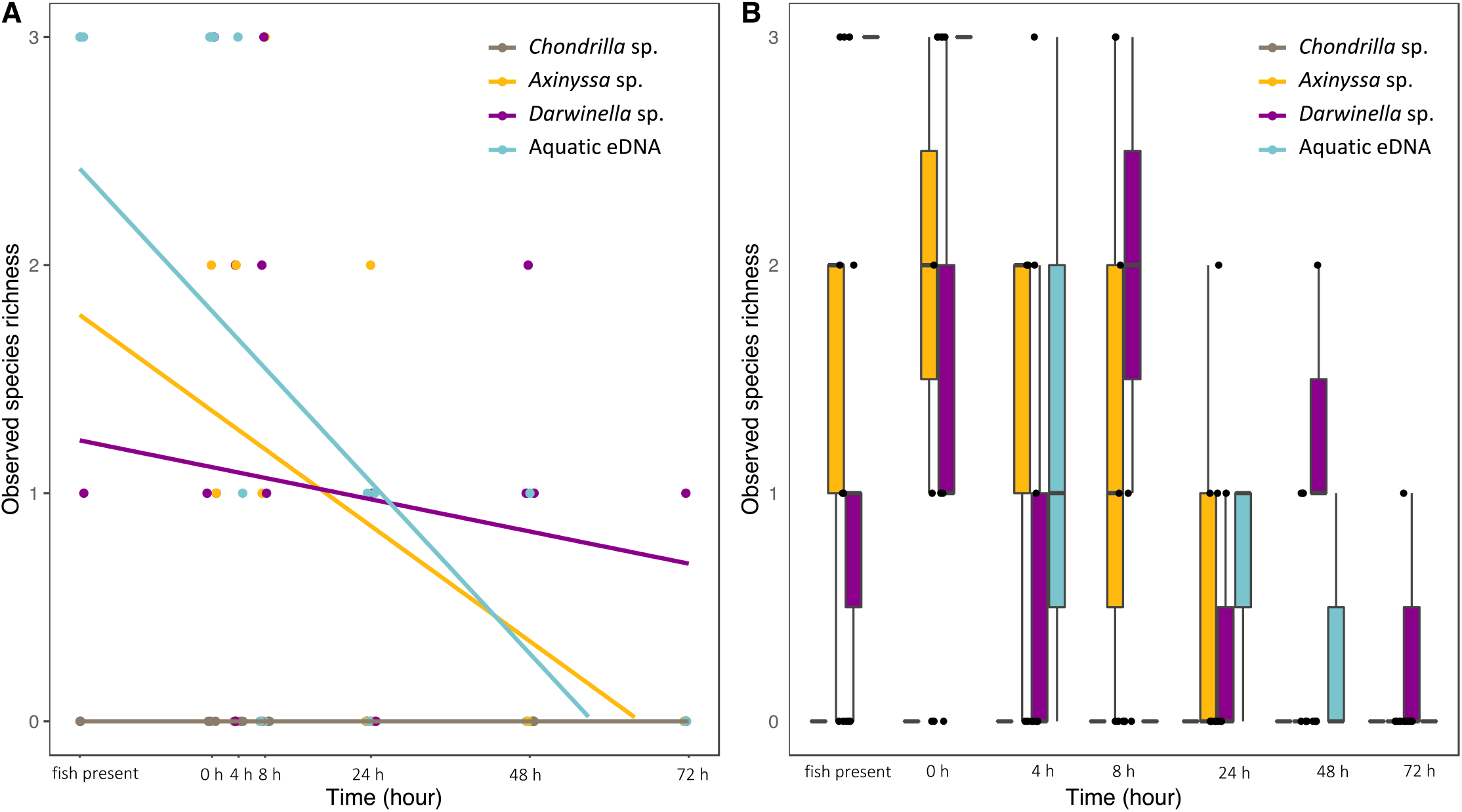
Comparison of alpha diversity among sample types. Read count data was converted to presence/absence data, thus, observed species richness was calculated using species incidence. The highest observed species richness is 3, indicating the sample can detect all fish. On the contrary, 0 indicates that no species are detected. **A**. The linear relationship between alpha diversity and time. **B**. Summary of species richness of each sample type at each time point. Colours and time code as in Fig. 2A.

Changes in fish community composition for aquatic eDNA and sponge nsDNA over time across tanks can be seen in Figure 4. The overall communities detected during the observation period did not significantly differ among sample types (P = 0.78, Df = 2), but community change was detected over time and more obviously in aquatic eDNA than in sponge nsDNA (Table S2). During the first 40 hours (when fish were present in the tanks), aquatic eDNA can consistently detect the entire community across the three tanks, while sponge nsDNA appeared to be less efficient in capturing the whole fish community (Fig. 4). However, after 4 hours of removing fish, eDNA degraded rapidly, and a drop in species detection with the aquatic eDNA was observed. On the contrary, fish community change was less notable in sponge nsDNA throughout the experimental period, especially in *Darwinella* sp. (P = 0.852, Df = 1). This result is consistent with the species richness pattern observed (see Fig. 3). Sponge nsDNA failed to detect the entire fish community in a single sampling event but detected fish over longer periods of time than the aquatic eDNA (Figs. 3, 4).

**Figure 4.**
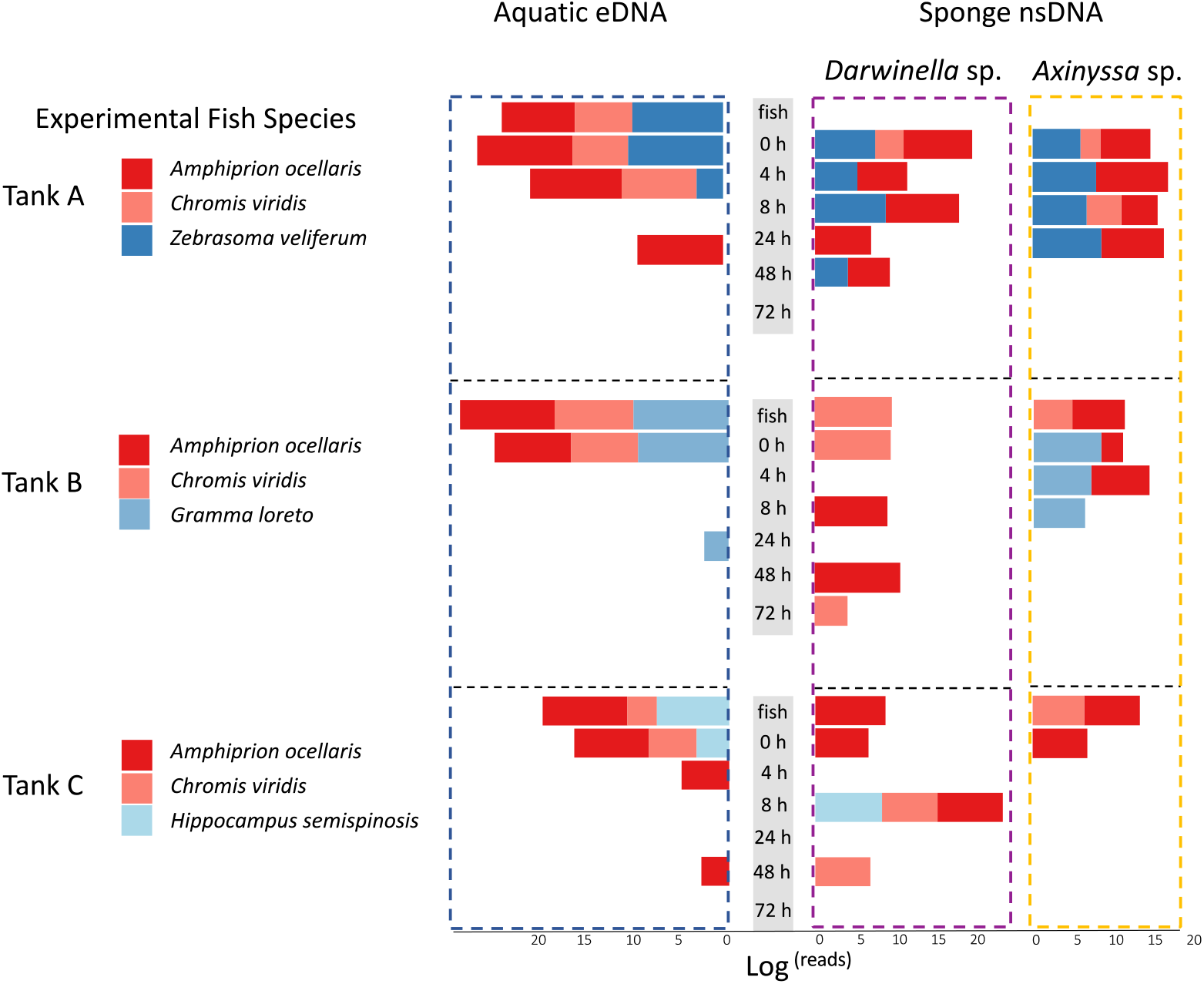
Community change over time. The horizontal panels represent three tanks; the vertical panels show three sample types. Water and sponge samples are separated by the timeline. For each tank, the community changes from top to bottom according to sampling time, and time codes as in Fig. 2A. The length of the bar is based on log_10_ read counts. When there is no bar at the corresponding time point, it represents that no fish species were detected. Colour codes for experimental fish species for each tank (Three species for each tank).

## Discussion

Our study confirms general expectations that sponges capture eDNA from the surrounding environment and emphasizes the variance in eDNA persistence and detectability among different sponge species. We employed three sponge species based on their established presence at the experimental facilities, and each one performed in a very distinct manner.

A key finding of this study was that fish species present in greater abundance or of larger size are more likely to be detected and contribute more reads to sponge nsDNA (Fig. 2B). This suggests that those sponges that are suitable for natural eDNA sampling might also be used to infer the relative abundance of fish. Future studies in more diverse wild ecosystems should assess the degree to which nsDNA concentration is correlated with species abundance or size, although the use of metabarcoding read counts to estimate relative species abundance still requires much ground-truthing (Cristescu & Hebert, 2018). Despite the simplified fish community used in this study, without choosing any particular sponge species with well-studied morphology and filtration characteristics, it is reassuring to see that at least some sponges are comparable with aquatic eDNA filtration, which is a powerful tool for monitoring fish population densities (Levi et al., 2019; Di Muri et al., 2020), leading to the expectation that sponges could indeed be adopted for such applications in the future. Further sponge nsDNA-based studies could be carried out also to examine the quantitative methods developed for aquatic eDNA, such as qPCR and ddPCR based on species-specific primers (Baker et al., 2018; Levi et al., 2019).

Another primary goal of this study was to investigate eDNA decay in sponges compared to water samples. Our results suggest that eDNA decay rates differ in sponge nsDNA and aquatic eDNA samples. For aquatic eDNA, as soon as the fish were removed from the tanks, the eDNA degraded rapidly over a 4-hour period. After that, fish species were poorly detected in the aquatic eDNA samples (Fig. 4). In contrast, fish can be detected for longer periods in sponge nsDNA samples; especially in *Darwinella* sp., eDNA did not degrade dramatically over time – at least over the short time frame of the experiment (72 hours). This is likely because these sponges are moderately effective pumpers, trapping eDNA efficiently and aided by a slow metabolism, allowing them to preserve eDNA over longer periods (Moitinho-Silva et al., 2017). In the case of delayed detection of very low eDNA concentrations, it might take time for a sponge to accumulate eDNA in its tissue to a detectable level. However, due to the interaction between sponge physiology and symbiont content (Ribes et al., 1999; Hentschel et al., 2006; Leys et al., 2012), it is possible that some sponges may have faster eDNA decay rates than aquatic eDNA in open water. Here our empirical evidence shows that some sponges will preserve DNA for longer than it persists in water. It is possible to see how this feature could also be beneficial in monitoring rare migratory species from remote areas.

As demonstrated by previous studies (Mariani et al., 2019; Turon et al., 2020), sponges were able to detect pelagic or migratory species visiting some locations infrequently. Our evidence further demonstrates how certain sponges could help monitor these more elusive species (preserving eDNA for longer), which may be missed by other monitoring methods. As for the resident species, sponge nsDNA may provide insightful fish detection compared to aquatic eDNA because spatial and temporal variability of aquatic eDNA may affect species detection (Allan et al., 2021; Canals et al., 2021). Thus, the sessile nature of sponges could more exhaustively track fish’s diel fluctuations and other behaviours. Furthermore, several sponge species can typically settle on various human-made structures (e.g., piers, moorings, and oil rigs) and colonise artificial reefs. Therefore, sponges are ideal candidates for linking biodiversity assessment with human impact (Wulff et al., 2001; Vad et al., 2021). Although the natural sampler approach is in its infancy and requires much validation in natural scenarios, it can be hypothesized that the many factors known to promote degradation and transport of aquatic eDNA (Strickler et al., 2015; Collins et al., 2018; Holman et al., 2021) could be less influential on sponges, given that eDNA would be afforded protection inside their tissue.

Nevertheless, biotic processes, mostly linked to morphology, filtration rate, physiology, and symbiosis, necessitate further research. In our study, *Chondrilla* sp. failed to detect any fish. This could be due to the intrinsic inability of this species to capture or preserve eDNA. Members of the genus *Chondrilla* are high microbial abundance (HMA) sponges, compared to their counterparts of the genus *Axinyssa* and *Darwinella* (Moitinho-Silva et al., 2017; Batista et al., 2018; Díez-Vives et al., 2020). HMA sponges contain more abundant symbiont communities and content that could slow filtration and accelerate metabolism, thus, influencing the capture and persistence of eDNA. HMA sponges filter 30–40% slower than low microbial abundance (LMA) sponges (Weisz et al., 2007), even though chondrosiids are relatively effective pumpers (Milanese et al., 2003). Therefore, *Chondrilla* sp. is a relatively disadvantaged sponge in the competition for eDNA capture in tanks, compared to the LMA sponges (*Axinyssa* sp. and *Darwinella* sp.). Furthermore, exposure to manipulative stress, such as those in our experiment, may also affect its filtration characteristics, and PCR inhibitors, such as pigments and natural products, or even an aggressive microbiome, may all play a part in eDNA detection from sponge tissue. Therefore, collecting several sponge species from the same habitat could serve as a good strategy for improving fish community detection in natural settings. A previous study has also shown that not all sponges have the same ability to capture and retain eDNA, yet the sponge morphology did not significantly affect the detected OTU richness (Turon et al., 2020). We also note that a single sponge sample may not be sufficient to represent an entire fish community in our experiment. Overall, a deeper understanding of the underlying mechanism of how sponges capture and preserve eDNA will be helpful for users to design different applications.

## Conclusion

While we are still far from a standardised workflow for sponge nsDNA, we can identify a number of directions towards that goal. Our study focused on comparing eDNA decay between sponge and water samples, but much work is still needed to optimize tissue preparation for DNA isolation, determine the size/quantity of sponge tissue required, evaluate the effects of various DNA extraction and PCR strategies that will influence eDNA detection, and biological and technical replication required for estimating biodiversity (Kumar et al., 2020; Bohmann et al., 2021). It will also be essential to ground-truth the performance of sponge nsDNA in multiple natural settings, using a variety of sponge species, in comparison with aquatic eDNA samples and established conventional methods. Once these outstanding challenges are met, sponge nsDNA will offer the advantages of a cost-effective method for the detection of fish diversity that is comparable to that of aquatic eDNA, which could significantly streamline sampling operations, reduce the use of plastic, and perhaps provide a number of biological features that will significantly enhance the toolkit of marine eDNA to further bolster biodiversity assessment.

## Acknowledgements

We are grateful to aquarium staff: Hartley George, Michelle Calvert and Matt Drysdale for helping us complete the tank setup and sampling. This study was supported by grant NE/T007028/1 from the UK Natural Environment Research Council. A.R. was also supported by the Spanish Ministry of Science and Innovation grant (RYC2018-024247-I) and CSIC intramural grant (202030E006).

## Statement of authorship

S.M., A.R. and L.R.H. designed the study, L.R.H., J.C., M.B.A. and E.F.N. handled specimens and collected samples, L.R.H. and E.F.N. performed the molecular experiments with assistance from P.S.; W.C. and P.S. performed the bioinformatic analyses. W.C. performed the statistical analyses and wrote the first draft of the manuscript, and all authors contributed substantially to interpretations and revisions.

## Data accessibility and Benefit-Sharing Statement

Raw sequencing data and all bioinformatics and statistical analysis files are deposited in the Zenodo. The archive is available for downloading at: https://doi.org/10.5281/zenodo.6331103.

All collaborators are included as co-authors.

## Reference

Aglieri, G., Baillie, C., Mariani, S., Cattano, C., Calò, A., Turco, G., Spatafora, D., Di Franco, A., Di Lorenzo, M., Guidetti, P., & Milazzo, M. (2021). Environmental DNA effectively captures functional diversity of coastal fish communities. Molecular Ecology, 30(13), 3127–3139. https://doi.org/10.1111/mec.15661

Allan, E. A., DiBenedetto, M. H., Lavery, A. C., Govindarajan, A. F., & Zhang, W. G. (2021). Modeling characterization of the vertical and temporal variability of environmental DNA in the mesopelagic ocean. Scientific Reports, 11(1), 1–15. https://doi.org/10.1038/s41598-021-00288-5

Andersen, K., Bird, K. L., Rasmussen, M., Haile, J., Breuning-Madsen, H., Kjær, K. H., Orlando, L., Gilbert, M. T. P., & Willerslev, E. (2012). Meta-barcoding of “dirt” DNA from soil reflects vertebrate biodiversity. Molecular Ecology, 21(8), 1966–1979. https://doi.org/10.1111/j.1365-294X.2011.05261.x

Baker, C. S., Steel, D., Nieukirk, S., & Klinck, H. (2018). Environmental DNA (eDNA) from the wake of the whales: Droplet digital PCR for detection and species identification. Frontiers in Marine Science, 5, 1–11. https://doi.org/10.3389/fmars.2018.00133

Batista, D., Costa, R., Carvalho, A. P., Batista, W. R., Rua, C. P. J., de Oliveira, L., Leomil, L., Fróes, A. M., Thompson, F. L., Coutinho, R., & Dobretsov, S. (2018). Environmental conditions affect activity and associated microorganisms of marine sponges. Marine Environmental Research, 142, 59–68. https://doi.org/10.1016/j.marenvres.2018.09.020

Bessey, C., Jarman, S. N., Berry, O., Olsen, Y. S., Bunce, M., Simpson, T., Power, M., McLaughlin, J., Edgar, G. J., & Keesing, J. (2020). Maximizing fish detection with eDNA metabarcoding. Environmental DNA, 2(4), 493–504. https://doi.org/10.1002/edn3.74

Bessey, C., Jarman, S. N., Simpson, T., Miller, H., Stewart, T., Keesing, J. K., & Berry, O. (2021a). Passive eDNA collection enhances aquatic biodiversity analysis. Communications Biology, 4(1). https://doi.org/10.1038/s42003-021-01760-8

Bessey, C., Gao, Y., Truong, Y. B., Miller, H., Jarman, S. N., & Berry, O. (2021b). Comparis on of materials for rapid passive collection of environmental DNA. Authorea, https://doi.org/10.22541/au.163707272.25577910/v1

Bohmann, K., Evans, A., Gilbert, M. T. P., Carvalho, G. R., Creer, S., Knapp, M., Yu, D. W., & de Bruyn, M. (2014). Environmental DNA for wildlife biology and biodiversity monit oring. Trends in Ecology & Evolution, 29(6), 358–367. https://doi.org/http://dx.doi.org/10.1016/j.tree.2014.04.003

Bohmann, K., Elbrecht, V., Carøe, C., Bista, I., Leese, F., Bunce, M., Yu, D. W., Seymour, M., Dumbrell, A. J., & Creer, S. (2021). Strategies for sample labelling and library preparation in DNA metabarcoding studies. Molecular Ecology Resources, 1–16. https://doi.org/10.1111/1755-0998.13512

Boussarie, G., Bakker, J., Wangensteen, O. S., Mariani, S., Bonnin, L., Juhel, J. B., Kiszka, J. J., Kulbicki, M., Manel, S., Robbins, W. D., Vigliola, L., & Mouillot, D. (2018). Environmental DNA illuminates the dark diversity of sharks. Science Advances, 4(5). https://doi.org/10.1126/sciadv.aap9661

Boyer, F., Mercier, C., Bonin, A., Le Bras, Y., Taberlet, P., & Coissac, E. (2016). obitools: A unix-inspired software package for DNA metabarcoding. Molecular Ecology Resources, 16(1), 176–182. https://doi.org/10.1111/1755-0998.12428

Canals, O., Mendibil, I., Santos, M., Irigoien, X., & Rodríguez-Ezpeleta, N. (2021). Vertical stratification of environmental DNA in the open ocean captures ecological patterns and behavior of deep-sea fishes. Limnology And Oceanography Letters, 6(6), 339–347. https://doi.org/10.1002/lol2.10213

Collins, R. A., Wangensteen, O. S., O’Gorman, E. J., Mariani, S., Sims, D. W., & Genner, M. J. (2018). Persistence of environmental DNA in marine systems. Communications Biology, 1(1), 1–11. https://doi.org/10.1038/s42003-018-0192-6

Cristescu, M. E. (2014). From barcoding single individuals to metabarcoding biological communities: Towards an integrative approach to the study of global biodiversity. Trends in Ecology and Evolution, 29(10), 566–571. https://doi.org/10.1016/j.tree.2014.08.001

Cristescu, M. E., & Hebert, P. D. N. (2018). Uses and misuses of environmental DNA in biodiversity science and conservation. Annual Review of Ecology, Evolution, and Systematics, 49, 209–230. https://doi.org/10.1146/annurev-ecolsys-110617-062306

Díez-Vives, C., Taboada, S., Leiva, C., Busch, K., Hentschel, U., & Riesgo, A. (2020). On the way to specificity - Microbiome reflects sponge genetic cluster primarily in highly structured populations. Molecular Ecology, 29(22), 4412–4427. https://doi.org/10.1111/mec.15635

Di Muri, C., Handley, L. L., Bean, C. W., Li, J., Peirson, G., Sellers, G. S., Walsh, K., Watson, H. V., Winfield, I. J., & Hänfling, B. (2020). Read counts from environmental DNA (eDNA) metabarcoding reflect fish abundance and biomass in drained ponds. Metabarcoding and Metagenomics, 4, 97–112. https://doi.org/10.3897/MBMG.4.56959

Eble, J. A., Daly-Engel, T. S., DiBattista, J. D., Koziol, A., & Gaither, M. R. (2020). Marine environmental DNA: Approaches, applications, and opportunities. Advances in Marine Biology, 86(1), 141–169. https://doi.org/10.1016/bs.amb.2020.01.001

Ereskovsky, A., Borisenko, I. E., Bolshakov, F. V., & Lavrov, A. I. (2021). Whole-body regeneration in sponges: Diversity, fine mechanisms, and future prospects. Genes, 12(4), 506. https://doi.org/10.3390/genes12040506

Ficetola, G. F., Pansu, J., Bonin, A., Coissac, E., Giguet-Covex, C., De Barba, M., Gielly, L., Lopes, C. M., Boyer, F., Pompanon, F., Rayé, G., & Taberlet, P. (2015). Replication levels, false presences and the estimation of the presence/absence from eDNA metabarcoding data. Molecular Ecology Resources, 15(3), 543–556. https://doi.org/10.1111/1755-0998.12338

Frøslev, T. G., Kjøller, R., Bruun, H. H., Ejrnæs, R., Brunbjerg, A. K., Pietroni, C., & Hansen, A. J. (2017). Algorithm for post-clustering curation of DNA amplicon data yields reliable biodiversity estimates. Nature Communications, 8, 1188. https://doi.org/10.1038/s41467-017-01312-x

Gerrodette, T., & Flechsig, A. O. (1979). Sediment-induced reduction in the pumping rate of the tropical sponge Verongia lacunosa. Marine Biology, 55, 103–110. doi: 10.1007/BF0039730

Goldberg, C. S., Turner, C. R., Deiner, K., Klymus, K. E., Thomsen, P. F., Murphy, M. A., Spear, S. F., McKee, A., Oyler-McCance, S. J., Cornman, R. S., Laramie, M. B., Mahon, A. R., Lance, R. F., Pilliod, D. S., Strickler, K. M., Waits, L. P., Fremier, A. K., Takahara, T., Herder, J. E., & Taberlet, P. (2016). Critical considerations for the application of environmental DNA methods to detect aquatic species. Methods in Ecology and Evolution, 7(11), 1299–1307. https://doi.org/10.1111/2041-210X.12595

Hansen, B. K., Bekkevold, D., Clausen, L. W., & Nielsen, E. E. (2018). The sceptical optimist: challenges and perspectives for the application of environmental DNA in marine fisheries. Fish and Fisheries, 19(5), 751–768. https://doi.org/10.1111/faf.12286

Hentschel, U., Usher, K. M., & Taylor, M. W. (2006). Marine sponges as microbial fermenters. FEMS Microbiology Ecology, 55(2), 167–177. https://doi.org/10.1111/j.1574-6941.2005.00046.x

Hoffmann, F., Røy, H., Bayer, K., Hentschel, U., Pfannkuchen, M., Brümmer, F., & De Beer, D. (2008). Oxygen dynamics and transport in the Mediterranean sponge Aplysina aerophoba. Marine Biology, 153(6), 1257–1264. https://doi.org/10.1007/s00227-008-0905-3

Holman, L. E., Chng, Y., & Rius, M. (2021). How does eDNA decay affect metabarcoding experiments? Environmental DNA, 1–9. https://doi.org/10.1002/edn3.201

Kahn, A. S., Yahel, G., Chu, J. W. F., Tunnicliffe, V., & Leys, S. P. (2015). Benthic grazing and carbon sequestration by deep-water glass sponge reefs. Limnology and Oceanography, 60(1), 78–88. https://doi.org/10.1002/lno.10002

Kandler, N. M., Wooster, M. K., Leray, M., Knowlton, N., de Voogd, N. J., Paulay, G., & Berumen, M. L. (2019). Hyperdiverse macrofauna communities associated with a common sponge, stylissa carteri, shift across ecological gradients in the central red sea. Diversity, 11(2), 1–16. https://doi.org/10.3390/d11020018

Kirtane, A., Atkinson, J. D., & Sassoubre, L. (2020). Design and Validation of Passive Environmental DNA Samplers Using Granular Activated Carbon and Montmorillonite Clay. Environmental Science and Technology, 54(19), 11961–11970. https://doi.org/10.1021/acs.est.0c01863

Kolde, R. (2019). Pheatmap: pretty heatmaps. R Package Version 1.0.12.

Kumar, G., Eble, J. E., & Gaither, M. R. (2020). A practical guide to sample preservation and pre-PCR processing of aquatic environmental DNA. Molecular Ecology Resources, 20(1), 29–39. https://doi.org/10.1111/1755-0998.13107

Lebuhn, G., Droege, S., Connor, E. F., Gemmill-Herren, B., Potts, S. G., Minckley, R. L., Griswold, T., Jean, R., Kula, E., Roubik, D. W., Cane, J., Wright, K. W., Frankie, G., & Parker, F. (2013). Detecting Insect Pollinator Declines on Regional and Global Scales. Conservation Biology, 27(1), 113–120. https://doi.org/10.1111/j.1523-1739.2012.01962.x

Levi, T., Allen, J. M., Bell, D., Joyce, J., Russell, J. R., Tallmon, D. A., Vulstek, S. C., Yang, C., & Yu, D. W. (2019). Environmental DNA for the enumeration and management of Pacific salmon. Molecular Ecology Resources, 19(3), 597–608. https://doi.org/10.1111/1755-0998.12987

Leys, S. P., & Hill, A. (2012). The Physiology and Molecular Biology of Sponge Tissues. Advances in Marine Biology, 62, 1–56. https://doi.org/10.1016/B978-0-12-394283-8.00001-1

Lynggaard, C., Bertelsen, M. F., Jensen, C. V., Johnson, M. S., Frøslev, T. G., Olsen, M. T., & Bohmann, K. (2022). Airborne environmental DNA for terrestrial vertebrate community monitoring. Current Biology, 1–7. https://doi.org/10.1016/j.cub.2021.12.014

Mahé, F., Rognes, T., Quince, C., de Vargas, C., & Dunthorn, M. (2015). Swarmv2: Highly-scalable and high-resolution amplicon clustering. PeerJ, 2015(12), 1–12. https://doi.org/10.7717/peerj.1420

Mariani, S., Baillie, C., Colosimo, G., & Riesgo, A. (2019). Sponges as natural environmental DNA samplers. Current Biology, 29(11), R401–R402. https://doi.org/10.1016/j.cub.2019.04.031

McQuillan, J. S., & Robidart, J. C. (2017). Molecular-biological sensing in aquatic environments: recent developments and emerging capabilities. Current Opinion in Biotechnology, 45, 43–50. https://doi.org/10.1016/j.copbio.2016.11.022

Milanese, M., Chelossi, E., Manconi, R., Sarà, A., Sidri, M., & Pronzato, R. (2003). The marine sponge Chondrilla nucula Schmidt, 1862 as an elective candidate for bioremediation in integrated aquaculture. Biomolecular Engineering, 20(4–6), 363–368. https://doi.org/10.1016/S1389-0344(03)00052-2

Moitinho-Silva, L., Steinert, G., Nielsen, S., Hardoim, C. C. P., Wu, Y. C., McCormack, G. P., López-Legentil, S., Marchant, R., Webster, N., Thomas, T., & Hentschel, U. (2017). Predicting the HMA-LMA status in marine sponges by machine learning. Frontiers in Microbiology, 8, 1–14. https://doi.org/10.3389/fmicb.2017.00752

Morganti, T. M., Ribes, M., Yahel, G., & Coma, R. (2019). Size Is the Major Determinant of Pumping Rates in Marine Sponges. Frontiers in Physiology, 10, 1474. https://doi.org/10.3389/fphys.2019.01474

Pawlowski, J., Apothéloz-Perret-Gentil, L., & Altermatt, F. (2020). Environmental DNA: What’s behind the term? Clarifying the terminology and recommendations for its future use in biomonitoring. Molecular Ecology, 29(22), 4258–4264. https://doi.org/10.1111/mec.15643

Ribes, M., Coma, R., & Gili, J. M. (1999). Natural diet and grazing rate of the temperate sponge Dysidea avara (Demospongiae, Dendroceratida) throughout an annual cycle. Marine Ecology Progress Series, 176, 179–190. https://doi.org/10.3354/meps176179

Riesgo, A., Taboada, S., Kenny, N.J., Santodomingo, N., Moles, J., Leiva, C., Cox, E., Avila, C., Cardona, L. & Maldonado, M. (2021). Recycling resources: silica of diatom frustules as a source for spicule building in Antarctic siliceous demosponges. Zoological Journal of the Linnean Society, 192(2), 259–276. https://doi.org/10.1093/zoolinnean/zlaa058

Rognes, T., Flouri, T., Nichols, B., Quince, C., & Mahé, F. (2016). VSEARCH: A versatile open source tool for metagenomics. PeerJ, 4:e2584. https://doi.org/10.7717/peerj.2584

Ruppert, K. M., Kline, R. J., & Rahman, M. S. (2019). Past, present, and future perspectives of environmental DNA (eDNA) metabarcoding: A systematic review in methods, monitoring, and applications of global eDNA. Global Ecology and Conservation, 17, e00547. https://doi.org/10.1016/j.gecco.2019.e00547

Russo, T., Maiello, G., Talarico, L., Baillie, C., Colosimo, G., D’Andrea, L., Di Maio, F., Fiorentino, F., Franceschini, S., Garofalo, G., Scannella, D., Cataudella, S., & Mariani, S. (2021). All is fish that comes to the net: metabarcoding for rapid fisheries catch assessment. Ecological Applications, 31(2), 0–2. https://doi.org/10.1002/eap.2273

Schnell, I. B., Bohmann, K., & Gilbert, M. T. P. (2015). Tag jumps illuminated - reducing sequence-to-sample misidentifications in metabarcoding studies. Molecular Ecology Resources, 15(6), 1289–1303. https://doi.org/10.1111/1755-0998.12402

Sellers, G. S., Di Muri, C., Gómez, A., & Hänfling, B. (2018). Mu-DNA: A modular universal DNA extraction method adaptable for a wide range of sample types. Metabarcoding and Metagenomics, 2, 1–11. https://doi.org/10.3897/mbmg.2.24556

Siegenthaler, A., Wangensteen, O. S., Soto, A. Z., Benvenuto, C., Corrigan, L., & Mariani, S. (2019). Metabarcoding of shrimp stomach content: Harnessing a natural sampler for fish biodiversity monitoring. Molecular Ecology Resources, 19(1), 206–220. https://doi.org/10.1111/1755-0998.12956

Stauffer, S., Jucker, M., Keggin, T., Marques, V., Andrello, M., Bessudo, S., Cheutin, M. C., Borrero-Pérez, G. H., Richards, E., Dejean, T., Hocdé, R., Juhel, J. B., Ladino, F., Letessier, T. B., Loiseau, N., Maire, E., Mouillot, D., Mutis Martinezguerra, M., Manel, S., … Waldock, C. (2021). How many replicates to accurately estimate fish biodiversity using environmental DNA on coral reefs? Ecology and Evolution, 11(21), 14630–14643. https://doi.org/10.1002/ece3.8150

Strickler, K. M., Fremier, A. K., & Goldberg, C. S. (2015). Quantifying effects of UV-B, temperature, and pH on eDNA degradation in aquatic microcosms. Biological Conservation, 183, 85–92. https://doi.org/10.1016/j.biocon.2014.11.038

Taberlet, P., Bonin, A., Zinger, L., & Coissac, E. (2018). Environmental DNA: For Biodiversity Research and Monitoring. Oxford University Press, Oxford. DOI: 10.1093/oso/9780198767220.001.0001.

Thomas, A. C., Howard, J., Nguyen, P. L., Seimon, T. A., & Goldberg, C. S. (2018). ANDe™: A fully integrated environmental DNA sampling system. Methods in Ecology and Evolution, 9(6), 1379–1385. https://doi.org/10.1111/2041-210X.12994

Turon, M., Angulo-Preckler, C., Antich, A., Præbel, K., & Wangensteen, O. S. (2020). More Than Expected From Old Sponge Samples: A Natural Sampler DNA Metabarcoding Assessment of Marine Fish Diversity in Nha Trang Bay (Vietnam). Frontiers in Marine Science, 7, 1–14. https://doi.org/10.3389/fmars.2020.605148

Vad, J., Barnhill, K. A., Kazanidis, G., & Roberts, J. M. (2021). Human impacts on deep-sea sponge grounds: Applying environmental omics to monitoring. Advances in Marine Biology, 89, 53–78. https://doi.org/10.1016/bs.amb.2021.08.004

Valdivia-Carrillo, T., Rocha-Olivares, A., Reyes-Bonilla, H., Domínguez-Contreras, J. F., & Munguia-Vega, A. (2021). Integrating eDNA metabarcoding and simultaneous underwater visual surveys to describe complex fish communities in a marine biodiversity hotspot. Molecular Ecology Resources, 21(5), 1558–1574. https://doi.org/10.1111/1755-0998.13375

Van Soest, R. W. M., Boury-Esnault, N., Vacelet, J., Dohrmann, M., Erpenbeck, D., de Voog d N. J., Santodomingo, N., Vanhoorne, B., Kelly, M., & Hooper, J. N. A. (2012). Global diversity of sponges (Porifera). PLoS ONE, 7(4), e35105. https://doi.org/10.1371/journal.pone.0035105

Wang, Y., Naumann, U., Wright, S. T., & Warton, D. I. (2012). Mvabund-an R package for model-based analysis of multivariate abundance data. Methods in Ecology and Evolution, 3(3), 471–474. https://doi.org/10.1111/j.2041-210X.2012.00190.x

Weisz, J. B., Lindquist, N., & Martens, C. S. (2008). Do associated microbial abundances impact marine demosponge pumping rates and tissue densities? Oecologia, 155(2), 367– 376. https://doi.org/10.1007/s00442-007-0910-0

Wells, C. D., Paulay, G., Nguyen, B. N., & Leray, M. (2021). DNA metabarcoding provides insights into the diverse diet of a dominant suspension feeder, the giant plumose anemone Metridium farcimen. Environmental DNA. https://doi.org/10.1002/edn3.225

Wulff, J. (2001). Assessing and monitoring coral reef sponges: Why and how? Bulletin of Marine Science, 69(2), 831–846.

